# Assay for Rapid Quantification of Capped and Tailed Intact mRNA

**DOI:** 10.1101/2025.04.02.646833

**Authors:** Rachel Y. Gao, Tianjing Hu, Amber W. Taylor, Randy Lacey, Keely N. Thomas, Caitlin McCormick, David F. Miller, Kathy L. Rowlen, Erica D. Dawson

**Author notes:** Address correspondence to Erica Dawson. VitriVax 5500 Central Ave, Boulder, CO 80301. 5836 SE Arcadia Rd, Shelton, WA 98584. Corgenix Medical Corporation 11575 Main St #400, Broomfield, CO 80020.

## Abstract

The rapid development of mRNA vaccines since the COVID-19 pandemic has highlighted the potential of mRNA technology for vaccine and therapeutic applications. However, evaluating mRNA quality, including its integrity, 5′ capping efficiency, and 3′ poly(A) tailing, remains complex and challenging. Current analytical methods provide detailed insights, but are often limited by long analysis times, complex sample preparation, and the need for specialized expertise. The 5’ Cap***Q*** assay addresses these challenges and provides a novel, rapid, and straightforward method that offers a single, unique measurement of intact mRNA with both a 5′cap and 3′poly(A) tail. This 2-hour benchtop-based assay provides quantitative or relative analysis, with high accuracy and precision without extensive sample processing. The 5’ Cap***Q*** assay offers a significant advancement in mRNA analytics to alleviate bottlenecks in applications including *in vitro* transcription (IVT) reaction optimization, post-transcriptional capping optimization, and assessment of batch-to-batch IVT bioprocess consistency.

## 1. Introduction

The rapid development of mRNA vaccines in response to the 2020 global COVID-19 pandemic demonstrated that mRNA vaccines are a potent alternative to traditional vaccines, with the capacity for rapid clinical advancement.[1–15] Since the first two mRNA vaccines were licensed in December 2020, there are now mRNA vaccines in late-stage development aimed at preventing diseases such as tuberculosis, malaria, influenza, COVID-19, respiratory syncytial virus (RSV), Lyme disease and others.[1, 14, 16–19] In addition, mRNA technology has expanded well beyond vaccines and is being explored for various therapeutic applications, including immunotherapies, protein replacement, and genome editing,[14, 16, 18, 19] further supporting that mRNA technologies will continue to be important to global health.

The urgent need to develop COVID-19 vaccines to curb the pandemic led to a reliance on existing mRNA analytical methodologies, which were quickly adapted out of necessity.[13, 20–22] Many of the methods currently utilized and outlined below are not at-line measurements and are typically performed at centralized analytical labs or outsourced to CDMOs. These existing methods often require execution by scientists with specialized training, and sample analysis is sometimes further delayed by long wait times in queues. Currently, manufacturers perform up to three separate measurements to assess mRNA integrity, 5′ capping efficiency, and 3′ poly(A) tailing that collectively provide information regarding the overall intactness of the mRNA from 5′ cap to 3′ poly(A) tail.[23]

As outlined in the recent third draft USP guidance document for mRNA vaccine analytics,[23] mRNA integrity is typically evaluated using capillary electrophoresis (CE), capillary gel electrophoresis (CGE), and agarose gel electrophoresis (AGE).[24–27] CE is a technique that separates ions based on their size-to-charge ratio within a narrow capillary tube under the influence of an electric field, offering high efficiency and resolution.[24–27] CGE is a variant of CE, where the capillary is filled with a gel matrix, allowing the separation of mRNA based on size.[24–27] AGE is a widely used method for analyzing nucleic acids that separates RNA fragments by applying an electric field to a gel matrix made of agarose.[24] CE and CGE provide high-resolution separation of RNA fragments with fluorescence detection, making them highly precise once optimized, but these techniques often require extensive upfront condition optimization.[24, 25, 28] AGE is simpler and more accessible, but has lower resolution and relies on subjective visual analysis, making it less ideal for detecting subtle RNA degradation.[24] It is important to note that none of these electrophoretic methods provide information regarding the presence of the 5′ cap.

Per USP draft guidelines, 5′ capping efficiency also must be assessed, typically using reverse-phase liquid chromatography mass spectrometry (RP–LC-MS), ion pair reversed-phase high-performance liquid chromatography (IP-RP-HPLC), or liquid chromatography mass spectrometry (LC-MS).[23, 24, 29] RP-LC-MS provides high resolution detection of 5′ cap structures, IP-RP-HPLC enhances separation of nucleic acids through hydrophobic interactions with ion-pairing agents, and LC-MS enables precise identification and quantification of modified RNA components.[24] All three methods require some type of RNA fragmentation prior to analysis, typically via an enzymatic digestion step.[24] All three offer highly precise quantification of capped and uncapped mRNA but require lengthy and complex sample preparation, and require assay optimization to ensure accurate quantification.[24] Similar to 5′ capping efficiency, 3′ poly(A) tailing is assessed using IP-RP-HPLC and LC-MS and requires upfront enzymatic digestion or fragmentation of mRNA.[24] The current utilized methods provide rich characterization of mRNA constructs including insights into mRNA fragmentation and degradation pathways and poly(A) tail length dispersity. Certainly, these methods will continue to provide utility moving forward throughout an mRNA vaccine or therapeutic development program. As previously mentioned, bottlenecks exist, including often lengthy analysis time, the requirement for up front sample processing, the need for specialized expertise to execute the methods, and an overall requirement to execute multiple methods to get a full picture of how much fully intact, cap to tail, mRNA is present. Given these shortcomings and that mRNA should be intact with both a cap and tail for proper translation to functional protein *in vivo*, we undertook development of a rapid, straightforward method that can be executed at the benchtop in the bioprocess lab to alleviate these bottlenecks and facilitate bioprocess development.

In contrast to current methods, the method described herein called 5′ Cap***Q*** provides a single unique measurement of intact mRNA with both a 5′cap and 3′poly(A) tail and is designed for compatibility with InDevR’s VaxArray platform. VaxArray has been historically used for antigen quantification in traditional protein-based vaccines and serum antibody quantification,[30–36] and is currently in use by a wide variety of vaccine developers and manufacturers worldwide. The 5’ Cap***Q*** measurement can be performed in a relative manner for comparisons of different samples or preparations of the same mRNA construct, or quantitatively in which a sequence-matched standard of known integrity and capping efficiency (measured by orthogonal methods) is used as a calibration curve for quantification of unknown samples of interest. The VaxArray 5′ Cap***Q*** assay is rapid (less than 2 h), and does not require extensive sample preparation, assay optimization, or mRNA fragmentation or digestion. 5′ Cap***Q*** provides rapid results for % capped and intact mRNA with high accuracy and precision and is an additional mRNA analytical tool that should find high utility in applications such as *in vitro* transcription (IVT) reaction optimization, post-transcriptional capping optimization, and assessment of batch-to-batch IVT bioprocess consistency.

## 2. Materials and Methods

### 2.1 5’ Cap**Q** Assay Design

The 5′ Cap***Q*** assay is a microarray format immunoassay utilizing a monoclonal 5′ cap capture antibody to bind to 5′ cap structures on the mRNA construct of interest. In addition, a 20 nt PolyT oligonucleotide detection label modified with a fluorophore at both the 3’ and 5’ ends was designed to target the 3′poly(A) tail of the mRNA constructs. The assay can be produced in a 2×8 format with 16 wells per slide, as shown in **Figure 1a**, or in a 3×8 format as shown in **Figure 1b** with 24 wells per slide. The 5′ cap capture antibody was printed with 81 replicate spots in each well (**Figure 1c**) at InDevR using proprietary processes onto epoxide-functionalized glass using a piezoelectric microarray printing system.

**Figure 1.**
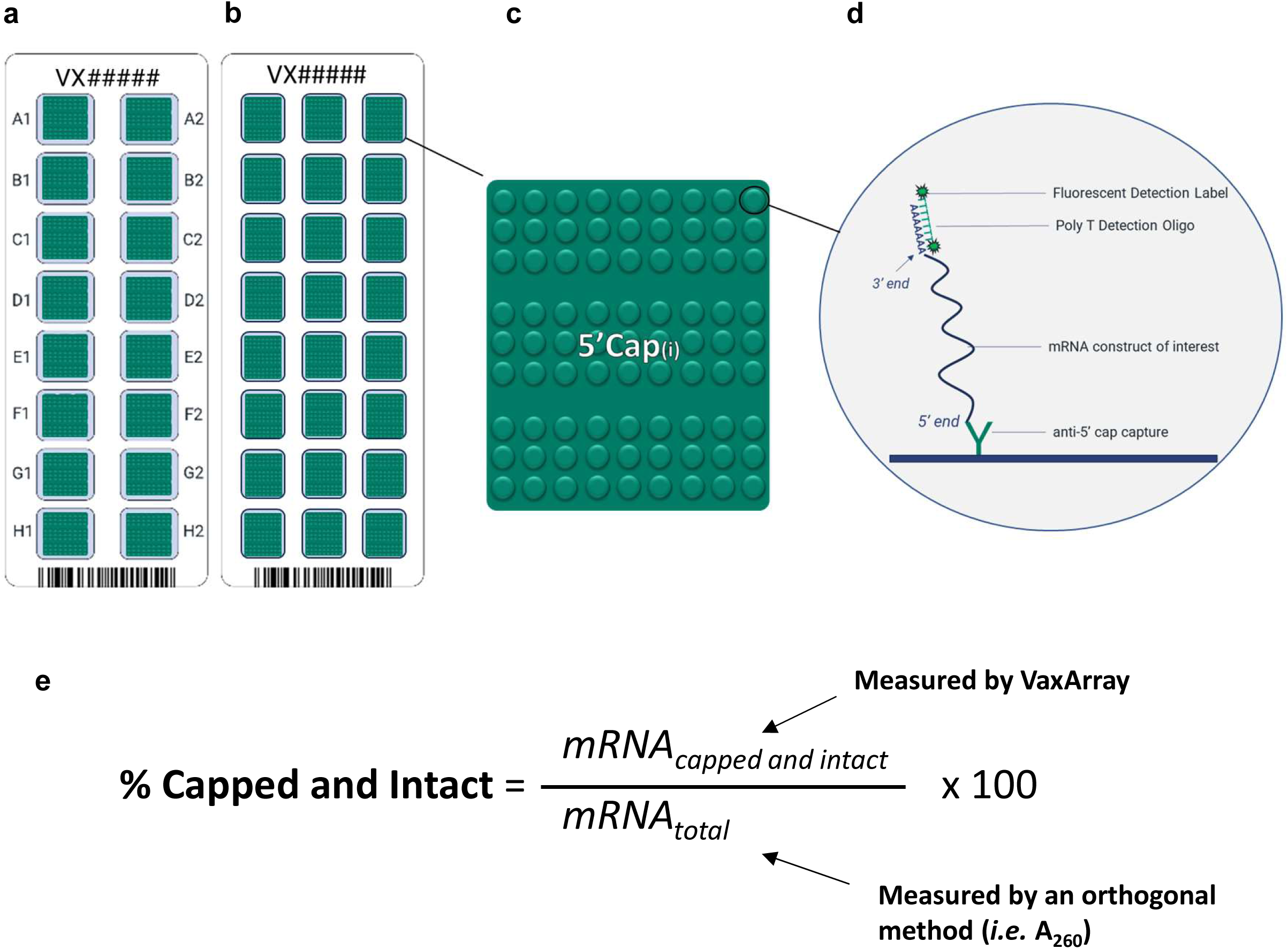
(a) Schematic of the VaxArray mRNA microarray slide showing 16 replicate microarrays or (b) 24 replicate microarrays, (c) individual microarray layout with 81 replicate spots of the printed anti-5’ cap antibody, and (d) assay detection scheme with capture via 5’ cap and PolyT labeling. (e) How VaxArray measures % capped and intact mRNA.

### 2.2 mRNA samples

Thirty-eight mRNA samples were obtained from six different sources, including both commercial suppliers and collaborators. Capped and uncapped mRNA materials were purchased from Aldevron (Fargo, ND) (source 1), Genscript BioTech (Piscataway, NJ) (source 4), and TriLink BioTechnologies (San Diego, CA) (source 6). Capped and uncapped mRNA materials were also obtained from collaborators at Biomanufacturing Training and Education Center (Raleigh, NC) (source 2), Cisterna Biologics (Carlsbad, CA) (source 3), and University of Pennsylvania (Philadelphia, PA) (source 5). Sequences for Mini HA and Mich NA were obtained from the literature[37] and synthesized by source 6.

mRNAs purchased from sources 4, 5, and 6 were ordered with one of several different codon optimization schemes applied, and the identifiers are appended with -1, -2, -3, or -4 denoting the scheme utilized for each construct. Codon optimization Scheme 1 followed a fundamental codon optimization method, Scheme 2 utilized GenScript Biotech’s method, Scheme 3 implemented GENEWIZ by Azenta (Burlington, MA) codon optimization, and Scheme 4 applied a modified fundamental codon optimization method. All mRNAs from sources 4 and 5 contain N1- methylpseudouridine-modified uridine (U), while source 6 is modified with 5-methoxy-U; all other sources contain a native uridine.

### 2.3 5’ Cap**Q** Assay Procedure

The assay follows a similar protocol to previously described VaxArray assays,[30–33] with the slide layout, microarray layout, and microarray hybridization and detection schemes depicted in **Figure 1a-d**. VaxArray slides were first equilibrated at 25⁰C for 30 minutes and placed inside a humidity chamber (VX-6204, InDevR Inc.) at room temperature. For the 2×8 format slide, 50 µL is added per well, whereas for the 3×8 format slide, 35 µL is added per well. Microarray slides were pre-washed with Wash Buffer A (VX-6324, InDevR Inc.) for 5 minutes. After the pre-wash, samples were diluted in an optimized 5′ Cap***Q*** mRNA Binding Buffer (2x) (VXI-6322, InDevR Inc.) at a final 1x concentration and applied to designated arrays. Slides were incubated for 1 hour and then washed with Wash Buffer A for 1 minute, followed by 5’ Cap***Q*** Poly T detection label (VXU- CAP-7601, InDevR Inc.) at a final concentration of 1x in an optimized 5′Cap***Q*** Detection Label Binding Buffer (2x) (VXI-6323, InDevR Inc.) for 30 minutes. The slides were then washed serially with Wash Buffer A, Wash Buffer B (VXI-6325, InDevR Inc.), and Wash Buffer C (VXI-6326, InDevR Inc.). Prior to drying the slides, excess liquid was pipetted off and the slides were placed in the ArrayDry (VXI-6218, VX-6220, InDevR Inc.) for a 30-second centrifugal drying step to remove any remaining liquid. Slides were imaged on the VaxArray Imaging System (VX-6000, InDevR Inc.), and downstream data analysis was performed using the VaxArray Analysis Software v3.1+.

For each sample and/or standard analyzed, a median fluorescence signal (RFU) value was calculated for each of the 81 spot locations across all replicate arrays for a given sample or standard to help account for array-to-array, slide-to-slide variation, and other sources of experimental variability. This analysis results in a set of 81 values representing the combined data over all replicates for each particular sample or standard, and each replicate analyzed is therefore not considered a separate technical replicate. Downstream statistics for each sample result presented herein such as mean, standard deviation, and confidence intervals were then performed on these 81 values.

### 2.4 Reactivity, Specificity, and LLOQ

Signal to background (S/B) ratios on the microarray were calculated to assess reactivity and specificity for 38 capped and 3 uncapped mRNA constructs from six sources, with a minimum reactivity threshold defined as a S/B of at least 3.0, and ideal specificity defined as a S/B of 1.0. Specificity of the 5′ cap capture antibody was verified by testing 10 nM uncapped mRNA constructs with Cap 0 and Cap 1 5′ caps. A label-only blank (no mRNA) was also analyzed to assess potential direct binding of the detection label to the anti-5′ cap antibody capture, with no direct binding was observed as evidenced by a S/B of 1.1 in the label-only blank. When comparing mRNA constructs obtained from different sources, all concentrations were converted to molar concentrations expressed in nM to account for variations in mRNA size and therefore enable more straightforward comparisons between samples.

The approximate LLOQ was determined by back-calculating the concentration at which the signal-to-background (S/B) ratio was equal to 3 using a linear regression to the linear portion of a 16- point curve.

### 2.5 Quantitative Analysis, Accuracy, and Precision

Quantitative % of capped and intact mRNA measurements were achieved by evaluating replicates of samples with unknown % capped and intact values against an average standard curve created from a sequence- and matrix-matched mRNA with a known concentration determined by A_260_, and ideally with known capping efficiency and integrity. The % capped and intact mRNA concentrations in the unknown samples were then back-calculated from the standard curve and divided by the total mRNA concentration measured by A_260_ (**Figure 1e**) and expressed as a percentage.

For the quantitation of 16 contrived samples (data in **Figure 4**, **Table 2**, and **Supplementary Table 2**), two users performed the experiment. An 8-pt dilution series of capped source 2 mRNA was analyzed starting from a maximum concentration of 0.7 μg/mL total mRNA. By mixing capped and uncapped sequence-matched mRNA, samples were contrived with the amount of capped mRNA therein ranging from 90% to 15% in 5% increments, all at a total mRNA concentration of 0.56 μg/mL. Each user processed four replicates of each contrived sample against the average of four standard curves distributed across four slides processed in the same experiment.

For quantitation of 2 contrived samples (data in **Figure 5**, **Table 3**, and **Supplementary Table 3**), three users each performed the following experiment twice (six experiments total). Two 8-pt dilution series of capped source 2 mRNA were analyzed starting from a maximum concentration of 0.32 μg/mL total mRNA. Two samples were contrived at 51% and 17% capped mRNA by mixing the capped and uncapped constructs. Each user processed eight replicates of each contrived sample against the average of two standard curves distributed across two slides processed in the same experiment.

Accuracy was assessed by comparing the measured % capped and intact mRNA to the expected values back-calculated from the calibration curve for the relevant experiment and expressed as a percentage of the expected result. Precision was expressed as percent relative standard deviation (% RSD), calculated by dividing the mean result by the standard deviation of the measurement and expressing as a percentage.

### 2.6 Relative Analysis

Relative % capped and intact mRNA is determined by evaluating samples with unknown amounts of % capped and intact mRNA at the same total mRNA concentration and comparing the signals generated (RFU) for each unknown sample.

For the relative analysis of eight samples (data in **Figure 6**, and **Table 4**), a single user processed four replicates of each contrived sample described below, with replicates distributed across 2 slides. Samples were contrived by mixing capped and uncapped sequence-matched mRNA to create samples ranging from 75% to 40% capped and intact mRNA in 5% increments, with all samples containing 0.86 μg/mL total mRNA.

## 3. Results and Discussion

### 3.1 Assay design

As described in the Methods section, an anti-5′ cap antibody was covalently immobilized to the microarray surface to enable capture of the 5′ cap structure, and a fluorescently labeled oligonucleotide targeting the poly(A) tail was used as a detection label, as schematically shown in **Figure 1**. Each microarray slide contains 16 or 24 replicate arrays with 81 replicate spots of the anti-5’ cap antibody printed in each array (**Figures 1a-c**). The anti-5′ cap antibody has been confirmed to capture both 5’ Cap 0 and Cap 1 structures. Because the detection label binds to the poly(A) tail on the 3′ end of the target mRNA construct, only mRNA containing a 5′ cap will be captured, and only mRNAs that also have a poly(A) tail will be labeled for detection as shown in **Figure 1d**.

Briefly, the assay (described in detail in the Methods section) takes less than 2 hours and consists of binding of the mRNA of interest in an optimized mRNA binding buffer to the anti-5’ cap antibody printed on the array, incubation of the detection label, a final wash, and fluorescence imaging and data analysis with the VaxArray Imaging System and software. To enable small differences in % capped and intact mRNA to be discerned, sample replicates and calibration curves are randomized on different arrays in the experiment to avoid sources of slide-to-slide or array-to-array systematic error.

The assay provides a single measurement of the total amount of intact mRNA in a sample that is both capped and tailed. This can be a relative measurement by comparing signals generated by samples comprised of the same construct (for example, as a function of differing IVT conditions, enzymatic capping conditions, or purification steps), or a quantitative measurement if measured alongside a sequence- and matrix-matched standard. As highlighted in **Figure 1e**, using the measurements of capping efficiency, intactness, and total mRNA concentration of the standard, % capped and intact mRNA of the unknown samples can be back-calculated.

### 3.2 Reactivity, Specificity, and Response with Dilution

Capped mRNA incorporating either Cap 0 or Cap 1 structures was sourced from six different sources as detailed in the Materials and Methods section, with corresponding sequence-matched uncapped mRNA obtained from three of the sources. All sources provided mRNA stock concentrations which were confirmed in-house by A_260_. **Figure 2a** displays the microarray layout, along with representative fluorescence images of Cap 0-containing mRNA, Cap 1-containing mRNA, and uncapped mRNA, from left to right. All were analyzed at 10 nM total mRNA. These images qualitatively indicate the microarray generates specific signals for mRNA with Cap 0 and Cap 1 structures and is not reactive with uncapped mRNA. **Table 1** further highlights that the assay is reactive to representative Cap 0 and/or Cap 1 mRNA from six different sources, as indicated by signal to background ratios (S/B) greater than 11.6 for all 6 constructs, and does not generate non-specific signal for uncapped mRNA from three different sources, as indicated by S/B ≤ 1.2 for all 3 uncapped constructs. While we did not focus on validating an analytical lower limit of quantification (LLOQ), given the intended applications of the assay are for bioprocess optimization and the concentrations of mRNA are likely to be well above the LLOQ, we did approximate the LLOQ for both a shorter 779 nt mRNA from source 2 at 0.34 nM, and for a longer 1,929 nt mRNA from source 6 at 0.62 nM. As detailed in the Materials and Methods section, these were approximated as the concentration at which the S/B exceeded 3 for a fit to a linear portion of a 16-point curve of each material.

**Figure 2.**
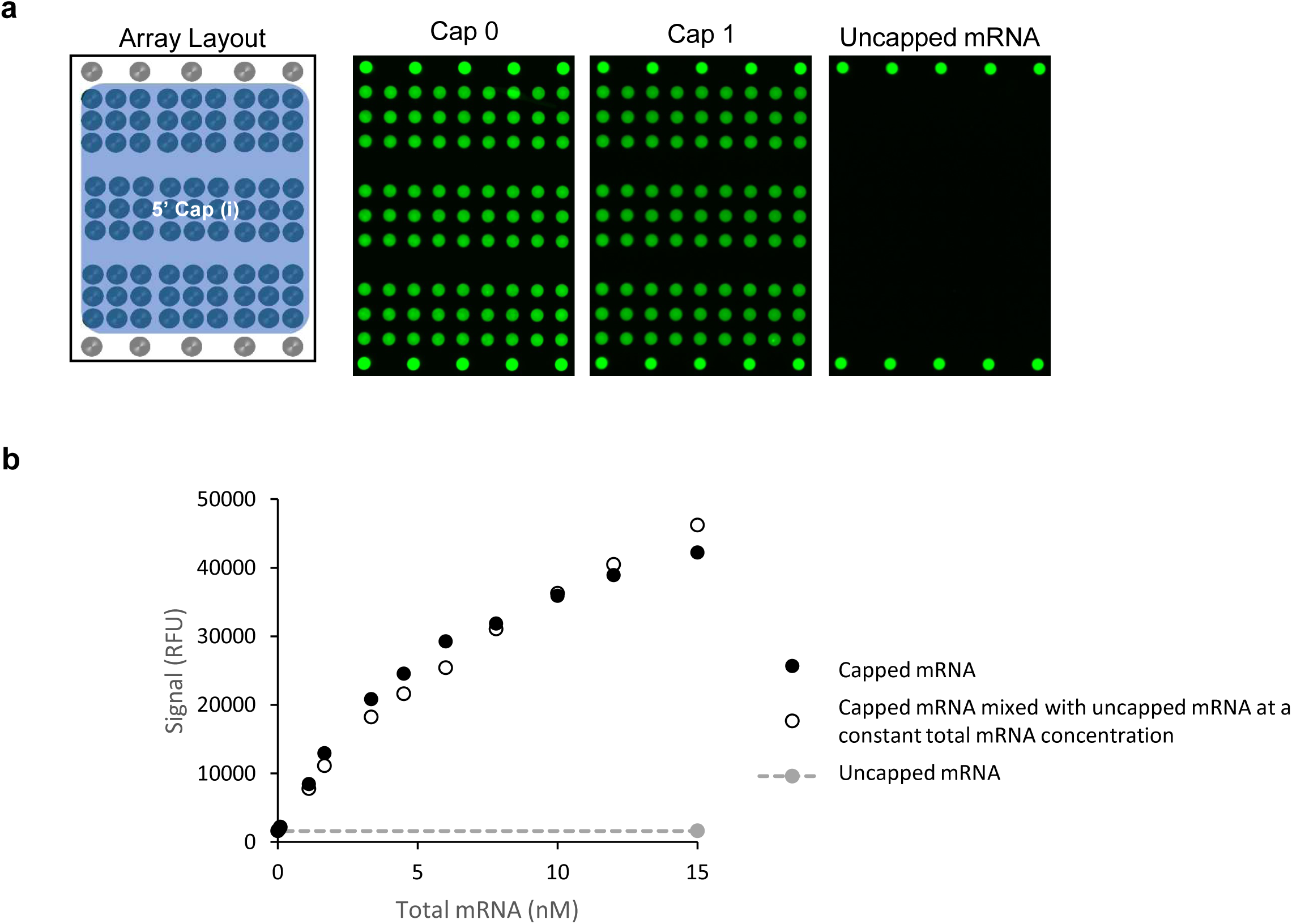
(a) Representative microarray images demonstrating the reactivity and specificity of 10 nM mRNA. Images from left to right correspond to Cap 0 mRNA (source 1), Cap 1 mRNA (source 2), and uncapped mRNA (source 1). (b) 16-point dilution curves of capped mRNA and capped mRNA mixed with uncapped mRNA at a constant total mRNA concentration, all Cap 0 (source 2).

**Table 1.** Reactivity and specificity of Cap 0 and Cap 1 constructs from six different suppliers.

| Source | Coding Region | Cap | Total Length (nt) | Poly(A) Tail Length (nt) | Signal to background (S/B) | Imaging exposure time (ms) |
| --- | --- | --- | --- | --- | --- | --- |
| 1 | Dasher green fluorescent protein (GFP) | Cap 1 | ~900 |  | **28.0** | 100 |
| 2 | Dasher GFP | Cap 0 | 779 |  | **27.8** | 200 |
| 3 | Red fluorescent protein (RFP) | Cap 0 | 799 | 40 | **19.5** | 500 |
| 4 | Influenza A/Wisconsin/67/2022 | Cap 1 | ~2052 |  | **11.6** | 1000 |
| 5 | Influenza A/Michigan/45/2015 | Cap 1 | ~2136 |  | **13.6** | 700 |
| 6 | Luciferase | Cap 1 | 1929 |  | **26.6** | 100 |
| 1 | Dasher GFP | None | ~900 |  | **1.1** | 1500 |
| 2 | Dasher GFP | None | 779 |  | **1.2** | 1500 |
| 3 | Red fluorescent protein (RFP) | None | 799 | 40 | **1.1** | 1500 |

Signal/background (S/B) for constructs (n=1 for each value).

To assess potential interference due to the presence of uncapped mRNA, **Figure 2b** compares 16-pt dilution curves of capped mRNA diluted in binding buffer to create the dilution series (black filled circles) and capped mRNA diluted with uncapped mRNA (from the same source) to keep a constant total mRNA concentration (open circles). Similar reactivity curves were observed for the sample contrived with uncapped mRNA and the dilution series of capped mRNA, indicating that no significant interference is observed due to the presence of uncapped mRNA. This indicates the ability to assess a sample containing a mixture of capped and uncapped material against a dilution series of a standard that may have a different concentration of capped and intact mRNA. Once again, the uncapped mRNA curve shows no reactivity up to 10 nM (indicated by grey filled circles).

**Figure 3** shows 4-point dilution series of the six capped mRNAs detailed in **Table 1**. The different mRNA constructs clearly have different reactivities and unique response curves, highlighted by imaging exposure times ranging from 100 ms to 1000 ms as noted in **Table 1**. An analysis of the mRNA features for the 6 constructs in Table 1 do not seem to indicate any obvious correlation between reactivity and features such as length, UTR sequences (each supplier utilizes unique 3’ and 5’ UTR sequences), coding region sequence, or cap structure. **Supplementary Table 1** includes additional reactivity data for all 38 unique capped mRNA constructs tested, with 33 constructs (87%) demonstrating reactivity. From the data in Supplemental Table 1, we cannot draw a clear correlation to any one specific variable as being the cause of the reactivity differences. Notably, all five unreactive mRNA constructs (S/B < 3) were obtained from source 4, and constructs from source 4 consistently exhibited lower signal intensity compared to mRNAs from other sources, with a maximum observed S/B of 11.6 at 1000 ms. All constructs from Source 4 share the same UTR, cap structure, and synthesis method, and some were reactive whereas others were not. We also note that each commercial mRNA source provided different levels and types of characterization data for the mRNAs received, and therefore we cannot rule out differences in mRNA integrity between suppliers or mRNA instability during storage as potential sources of reactivity differences. Different sources also contained varying uridine modifications, which may contribute to differences in reactivity. Some sources incorporated unmodified native uridine, while others included either N1-methyl-pseudouridine (source 4 and 5) or 5-methoxy-uridine (source 6) modifications. These differences in nucleotide composition could influence mRNA stability, translation efficiency, and immunogenicity [38, 39]. Lastly, it is possible that unique secondary or tertiary structures form with each mRNA construct and influence mRNA binding to the capture antibody or labeling efficiency of the polyT detection label.

**Figure 3.**
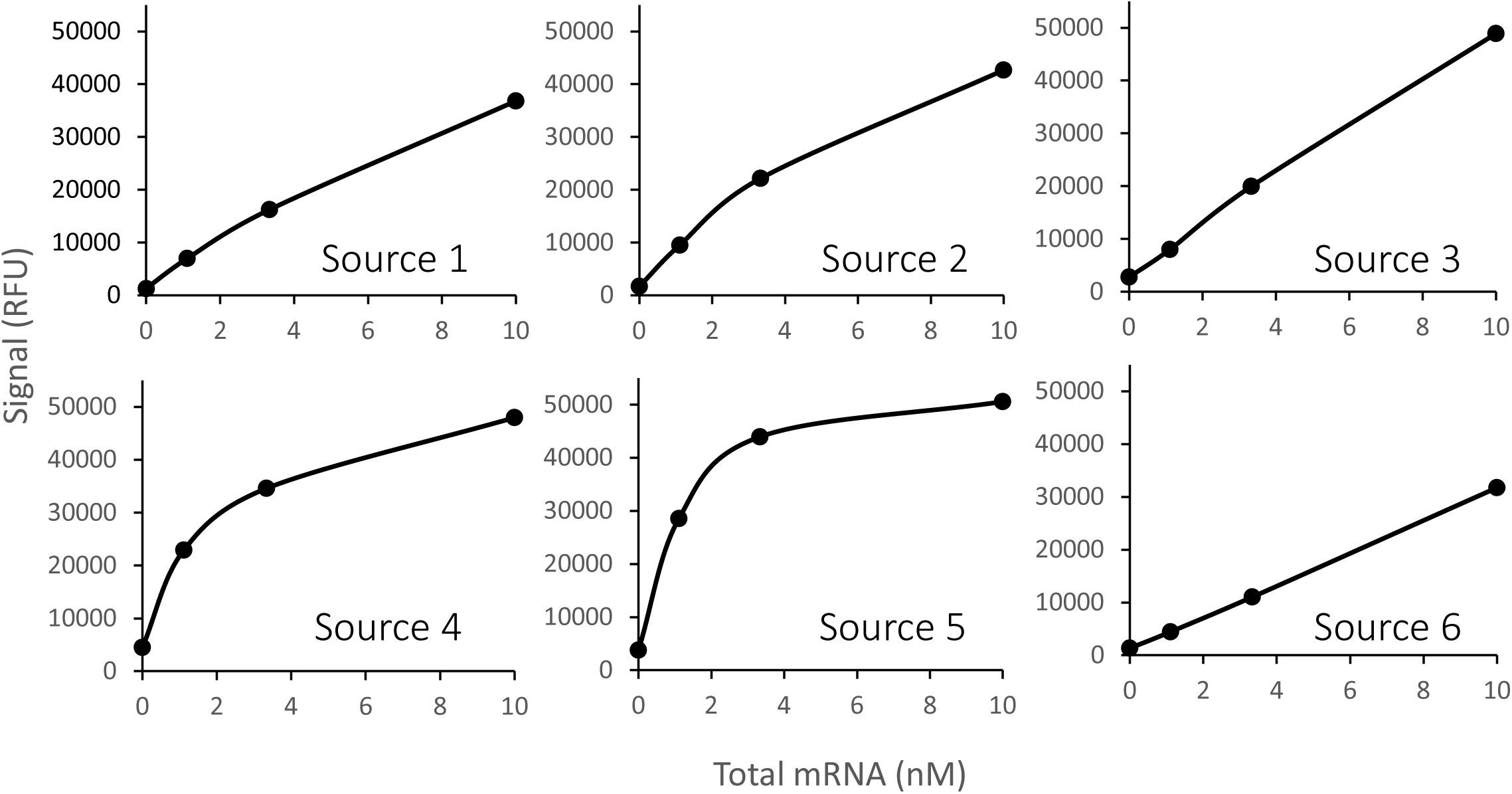
Representative response curves for mRNAs from 6 sources, with lines included for visual guidance only. 4-point curves starting at 10 nM. Imaging exposure times provided in Table 1.

These results demonstrate that the assay effectively detects capped mRNA from multiple sources, but that the reactivity varies between constructs, potentially due to differences in secondary and tertiary structures, uridine modifications, UTRs, poly(A) tail lengths, coding regions, size, and/or cap structures that result in differences in binding affinities for the capture antibody and/or detection label oligo. Several constructs were unreactive, but given they had the same cap structure as other constructs from the same provider that were reactive, we cannot attribute this to the cap structure itself. These reactivity differences clearly highlight the importance of an appropriate sequence-matched standard in any relative or quantitative analysis, and that relative signals between constructs that differ in sequence cannot be used as an indicator of the amount of capped and intact mRNA present. Therefore, the assay is best suited to applications in which a single construct is being optimized, for example in the optimization of *in vitro* transcription reactions to maximize the amount of intact and capped mRNA, optimizing post-transcriptional capping, or examining batch-to-batch consistency of *in vitro* transcription.

### 3.3 Accurate Quantification over a Wide Range of % Capped, Intact mRNA Values

To assess assay accuracy, we undertook a 4-slide quantitative analysis of 16 contrived samples prepared by mixing capped and uncapped mRNA with concentrations ranging from 90% to 15% capped and intact mRNA and assessed these against a dilution series of a standard curve prepared from the same capped starting material of known % capped and intact mRNA. This experiment was repeated by a second user.

**Figure 4a** shows the 8-point calibration curves for each user against which the unknowns were assessed. As can be seen from comparing the closed and open circles for users 1 and 2, respectively, both users obtained very similar responses for their calibration curves with excellent linearity with dilution as indicated by R^2^ >0.99. The concentration of % capped and intact mRNA in each of the 16 contrived samples for each user was back-calculated from the respective calibration curves, and **Figure 4b** shows the measured values obtained plotted against the expected values based on the sample preparation (data also shown in tabular format in **Table 2**). Overall, measured vs. expected % capped and intact mRNA in **Figure 4b** should be linear with slope=1. Both users obtained good linearity (User 1 R^2^=0.987, User 2 R^2^=0.991), indicating a high degree of correlation between expected and measured % capped and intact mRNA. In addition, the slope obtained for each user was close to 1, with a User 1 slope=0.934 and a User 2 slope=0.923. As shown in **Figure 4b** and **Table 2**, we also observe no overlap in the lower and upper 95% confidence limits (LCL and UCL) for samples that differ by 5% intact and capped mRNA indicating statistical confidence in measuring differences of 5%. As described in the Materials and Methods section, the statistical analysis approach data from each replicate microarray spot location across replicate arrays for each sample to account for potential array to array, slide to slide, and other experimental differences. With this experimental and analysis approach, small differences in % capped and intact mRNA can be distinguished with statistical confidence. Also shown in **Table 2**, accuracy ranged from 82.0% to 96.2% of the expected value for both operators and all samples, indicating that the assay is accurate over a wide range of % capped and intact mRNA values. **Supplementary Table 2** highlights the precision achieved for this experiment in terms of the %RSD of the signals for the n=81 spot locations pooled from the n=4 replicates run for each of the 16 contrived samples.

**Figure 4.**
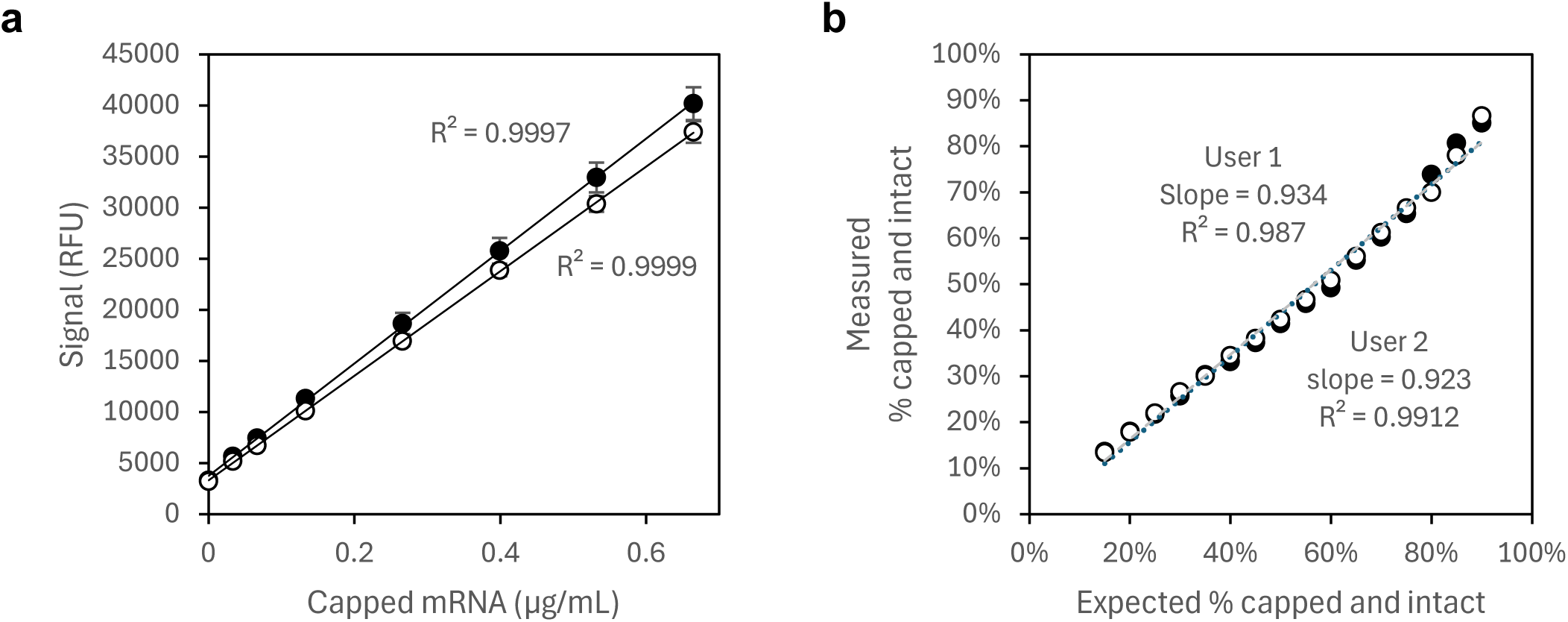
Quantitation of 16 contrived samples by User 1 (●) and User 2 (○). (a) 8-point dilution series of source 2 mRNA used as calibration curve, with error bars representing ±1 standard deviation of four replicate measurements at each concentration, and (b) back-calculated % capped and intact mRNA values from 2 users for 16 contrived samples, with error bars indicating the 95% confidence interval of n=81 median spot values from n=4 replicates. All fits shown are a single linear regression.

**Table 2.**
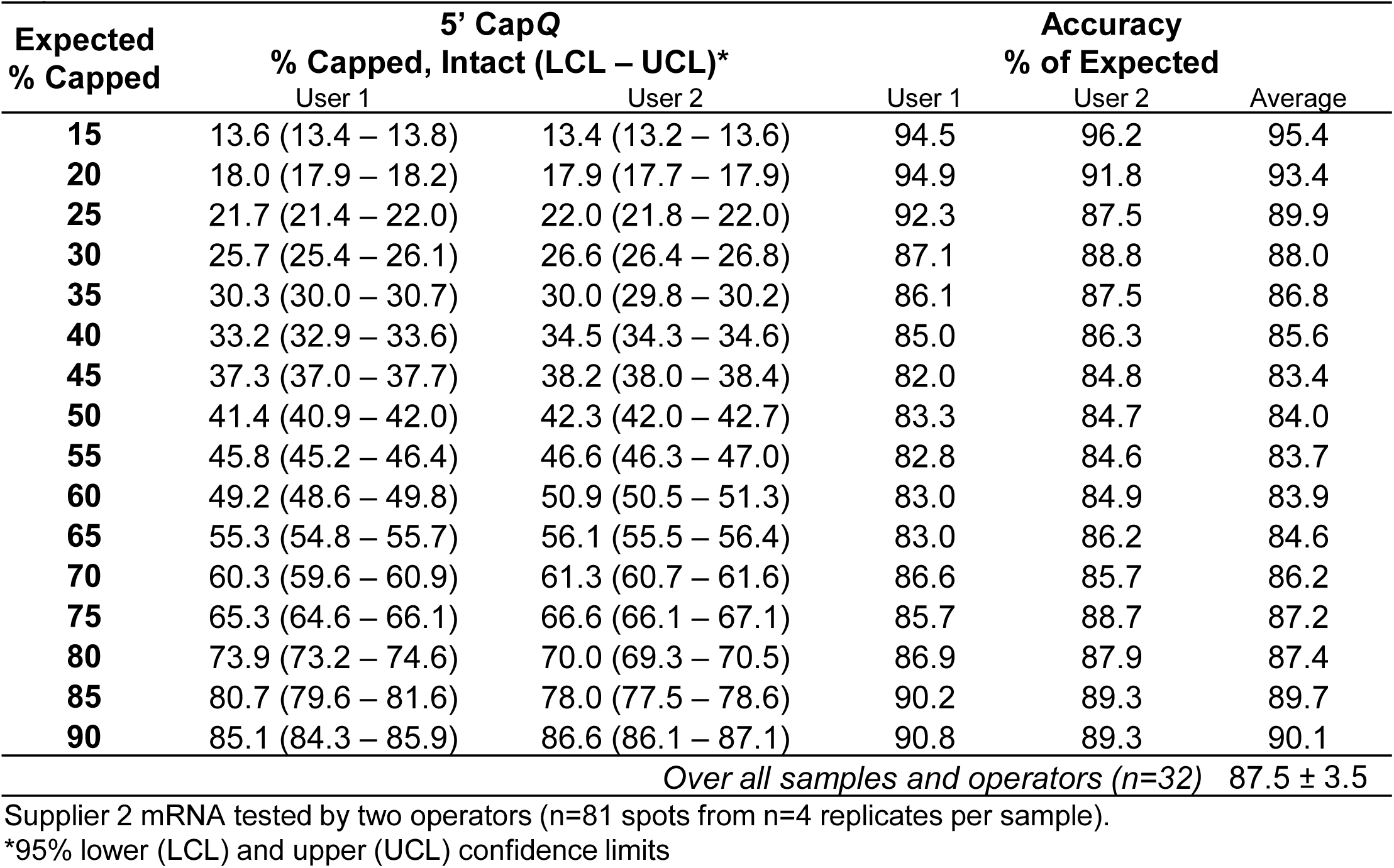
Accuracy and precision for comparison between users and across two different days in a 4-slide, 16-sample quantitative experiment.

These results demonstrate the assay’s quantification capabilities, with high reproducibility between users and a strong correlation between expected and measured % capped and intact mRNA. The observed linearity and accuracy indicate that the assay reliably quantifies capped mRNA across a wide range of concentrations of % capped and intact mRNA. Notably, the ability to distinguish samples differing by as little as 5% underscores its value in rapidly assessing mRNA integrity.

### 3.4 Precision and Accuracy Across Multiple Users and Days

To assess the precision and accuracy of the assay across multiple users and days, three operators each analyzed two contrived samples against a standard curve comprised of well-characterized mRNA of known % capped and intact concentration on two different days. For the contrived samples, two different mRNA mixtures were prepared at 51% and 17% capped and intact mRNA but at the same total mRNA concentration. **Figure 5** and **Supplementary Table 3** show the back-calculated % intact and capped mRNA values for the three users over each of two different testing days with the dotted line indicating the expected value for each of the two samples. **Table 3** shows the results of accuracy and precision in this multi-operator, multi-day study. Accuracy is shown as the % of the expected value (51% and 17%) for the contrived samples, and precision shown as the %RSD of n=81 spot locations pooled from 8 replicates. Across all operators, accuracy ranged from 84.1 to 116.8% of the expected result and precision ranged from 2.1 to 6.1 %RSD. These results demonstrate the high precision and accuracy of the assay across multiple operators and testing days indicating assay robustness. As previously mentioned, we also note that the use of a well-characterized, sequence-matched standard curve is critical to the assay’s ability to generate accurate measurements.

**Figure 5.**
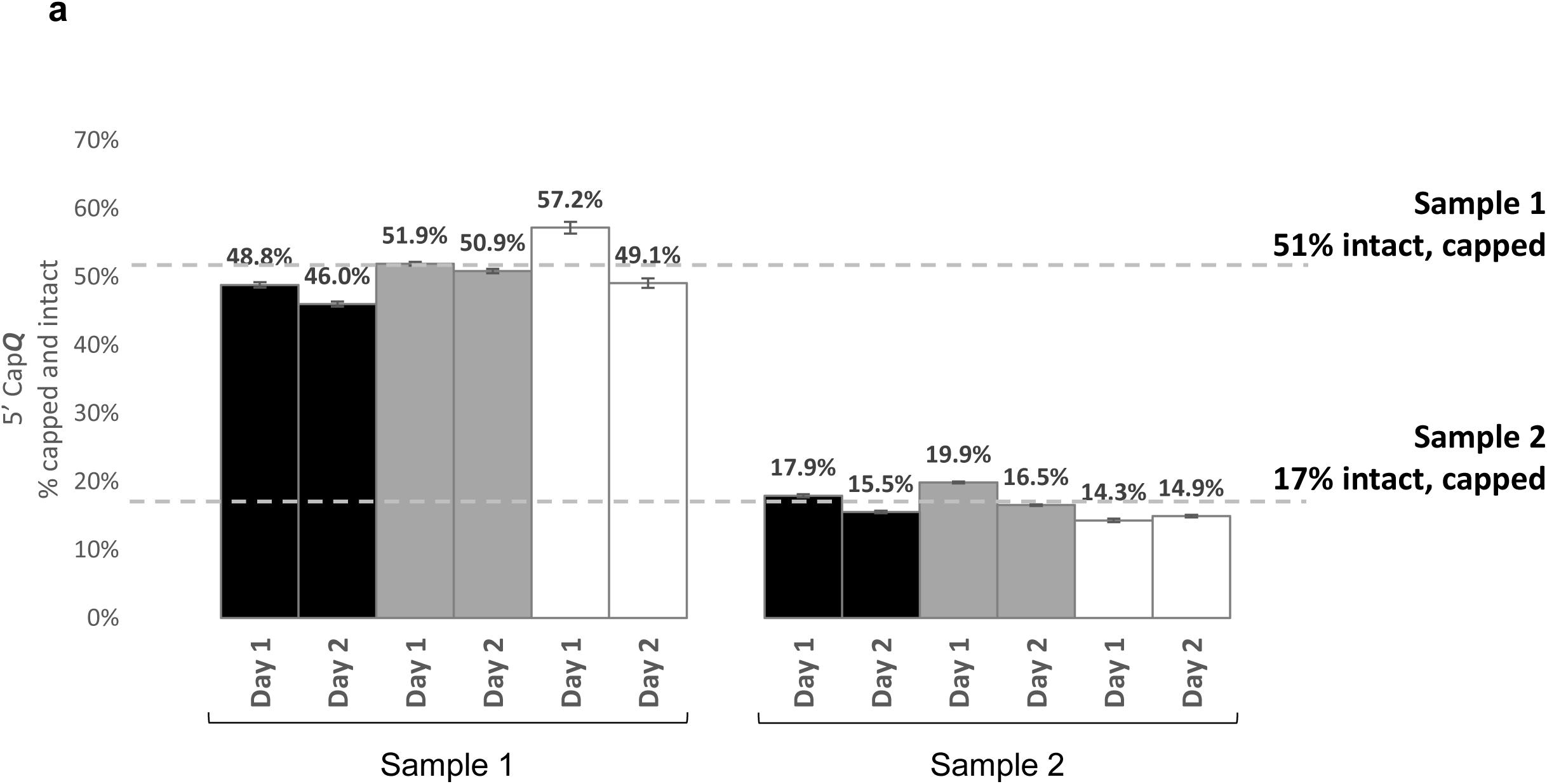
Day to day and user to user comparison for a 4-slide quantitative analysis. User 1 (black), User 2 (grey), and User 3 (white). Error bars indicate the 95% CI of n=81 median spot values from n=8 replicates.

**Table 3.**
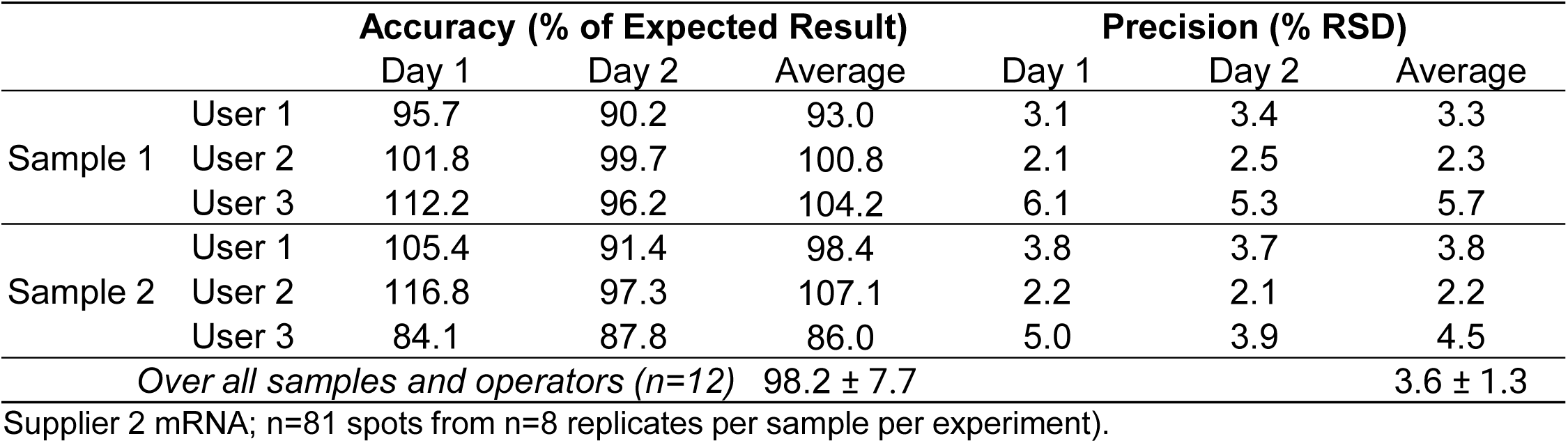
Accuracy and precision between three users and across two days in a 2-slide, 2-sample quantitative experiment.

### 3.5 Relative Analysis

If a well-characterized standard is not available, the assay can be used to perform a qualitative comparison based on differences in signal being correlated to differences in capped and intact mRNA. This is useful in scenarios where standards are not accessible or practical to generate, such as in early-stage experiments, routine screening, or when a relative comparison of materials is enough to inform optimization of reaction conditions or other experimental variables. To assess relative analysis, eight samples were prepared ranging from 75% to 50% capped and intact in 5% increments, with the total amount of mRNA remaining constant between all samples. **Figure 6** and **Table 4** display the median signal generated for each sample. Because there is no standard of known concentration, the samples should be prepared at the same total mRNA concentration (measured by a method like A_260_) such that differences in signal are not due to differences in total mRNA present, but only due to differences in the % of intact and capped mRNA. As shown in **Table 4**, we observe no overlap between the LCL and UCL (95%) of the signals for each sample, indicating significant differences between contrived samples. Precision was assessed by calculating the %RSD from 81 spots pooled across four replicates.

**Figure 6.**
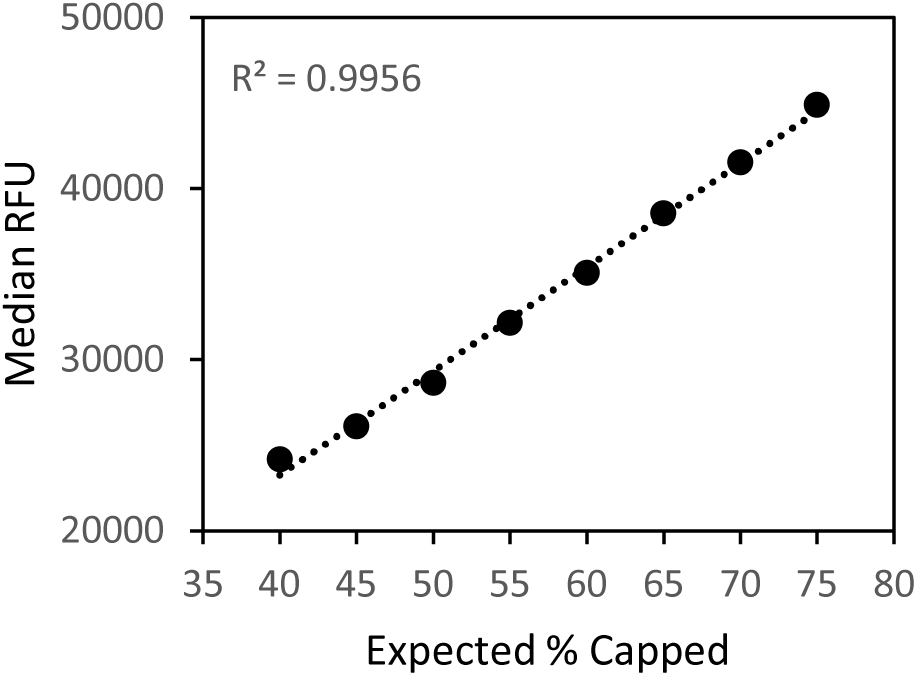
2-slide relative analysis of 8 contrived samples (R^2^ = 0.995, linear regression). Error bars indicate the 95% confidence interval (CI) of n=81 median spot values from n=4 replicates.

**Table 4.** Median RFU and 95% CI for a 2-slide relative analysis.

| **Expected**<br>**% Capped** | **Signal RFU**<br>**(LCL – UCL)\*** | **Precision**<br>**% RSD** |
| --- | --- | --- |
| **40** | 24170 (24010 - 24330) | 3.0 |
| **45** | 26090 (25910 - 26270) | 3.1 |
| **50** | 28630 (28390 - 28860) | 3.7 |
| **55** | 32140 (31910 - 32370) | 3.3 |
| **60** | 35060 (34790 - 35330) | 3.5 |
| **65** | 38540 (38270 - 38810) | 3.2 |
| **70** | 41540 (41230 - 41850) | 3.4 |
| **75** | 44890 (44580 - 45190) | 3.1 |
| *Over all samples (n=12)* |  | 3.3 ± 0.2 |

Supplier 2 mRNA (n=81 spots from n=4 replicates per sample per experiment).
\*Lower (LCL) and upper (UCL) 95% confidence limits

It is important to note that relative analysis is appropriate only when comparing samples with similar characteristics. Comparing constructs with different coding regions, UTRs, lengths, or cap structures would not yield valid conclusions in a relative analysis given that these factors can affect the reactivity of the construct. In addition, samples should be at least nominally purified or substantially diluted prior to analysis to minimize any potential matrix effects that could cause reactivity differences. Relative analysis offers valuable insights in specific contexts, and understanding its limitations helps ensure the most accurate and reliable results.

## 4. Conclusions

As the field of mRNA vaccines and therapeutics rapidly grows and diversifies, efficient and accurate characterization of mRNA 5′ capping, integrity, and 3′ poly(A) tailing is essential for ensuring the quality and consistency of the final vaccine product. While traditional methods such as CE, CGE, AGE, RP-LC-MS, IP-RP-HPLC, and LC-MS provide valuable insights into the characteristics of mRNA constructs,[23, 24, 29] they come with several challenges. These methods often require specialized training, expensive instrumentation, and labor-intensive procedures, which can be time-consuming and prone to variability. In addition, the necessity for mRNA fragmentation and method optimization can delay the analysis, hindering development timelines. An alternate method of mRNA characterization for use in bioprocess development should: (1) have excellent analytical performance, (2) deliver rapid results to enhance bioprocess efficiency, (3) demonstrate good specificity and reactivity, (4) have high ease of use to enable at-line analysis, (5) eliminate the need for specially trained personnel, and (6) have flexibility in throughput.

The introduction of the 5′ Cap***Q*** assay marks a significant addition in the field of mRNA characterization, offering a rapid and efficient method for assessing capped and intact mRNA without the need for complex sample preparation, fragmentation, or assay optimization often required with current methodologies. This assay utilizes an anti-5′ cap antibody to specifically capture the 5′ cap and labels the 3′ poly(A) tail, allowing for the capture and detection of only intact mRNA. By capturing and labeling opposite ends of the mRNA construct, it provides a unique measurement compared to existing methods. The 5’ Cap***Q*** assay is reactive with a wide range mRNA constructs and, importantly, uncapped mRNA does not interfere with reactivity of capped mRNA constructs (**Figure 2, 3**). When used quantitively, the assay provides precise and accurate measurements over a wide range of % capped and intact mRNA (**Table 2**), while also providing consistently accurate and precise data across users and days (**Figure 5**, **Table 3**).

The 5′ Cap***Q*** assay offers flexible throughput capabilities and a rapid time to result, allowing users to process and analyze a single slide (16 or 24 wells) or up to four slides (64 or 96 wells) in 2 hours. This speed and flexibility in sample throughput enables the generation of same-day results, helping to inform the bioprocess in near real-time. This is particularly beneficial for bioprocess applications where informed decisions and rapid turnaround times are crucial such as IVT reaction optimization, post-IVT enzymatic capping optimization, and assessing batch-to-batch IVT/bioprocess consistency. We are confident that the assay will find utility in a variety of bioprocess applications as a tool that can enhance the overall efficiency of mRNA vaccine and therapeutics production.

## Data Availability Statement

All relevant data from this study are available from the authors.

## Author Contributions

R.G. supervision, methodology, investigation, validation, formal analysis, visualization, writing-original draft, writing-review and editing; T.H. investigation, supervision, visualization, formal analysis, writing-review and editing; A.T. project administration, conceptualization, methodology, resources, visualization, writing-review and editing; R.L. investigation, formal analysis, visualization, writing-review and editing; K.T. methodology, investigation, writing-review and editing; C.M. methodology, investigation, writing-review and editing; D.M. supervision, resources, writing-review and editing; K.R. conceptualization, project administration, resources, writing-review and editing; E.D. conceptualization, project administration, supervision, methodology, resources, formal analysis, writing-original draft, writing-review and editing, visualization.

## Acknowledgements

We sincerely thank the laboratory of Jennifer Pancorbo, Biomanufacturing Training and Education Center (BTEC), NC State University, for providing capped and uncapped mRNA with sequences (source 2).

We sincerely thank Hari Bhaskaran, Cisterna Biologics, for providing capped and uncapped RFP mRNAs with sequences (source 3).

We sincerely thank the laboratory of Norbert Pardi, Department of Microbiology of the Perelman School of Medicine, University of Pennsylvania, for synthesizing capped mRNA and providing sequences (source 5).

## Supplementary Tables

**Supplementary Table 1.** Reactivity of 38 Capped mRNA Constructs

**Supplementary Table 2.** Accuracy and precision of two users on two different days in a 4 slide, 16-sample quantitative experiment

**Supplementary Table 3.** Expected % of capped and intact values for comparison between users and across two different days in a 2-slide, 2-sample quantitative experiment

