## Supplemental Table 1 for "Assay for Rapid Quantification of Capped and Tailed Intact mRNA"

**Supplementary Table 1.** Reactivity and Specificity of Capped mRNA constructs

| Source | Exposure (ms) | Construct | Signal to background ratio (S/B) |
|---|---|---|---|
| 1 | 100 | Dasher GFP | **28.0** |
| 2 | 200 | Dasher GFP | **27.8** |
| 3 | 500 | RFP | **19.5** |
| 4 | 1000 | Influenza A/Wisconsin/67/2022 – 2* | **11.6** |
| | | Influenza A/Wisconsin/67/2022 - 3 | **1.6** |
| | | Influenza A/Sydney/5/2021 - 1 | **1.7** |
| | | Influenza A/Wisconsin/588/2019 - 1 | **1.8** |
| | | Influenza A/Darwin/6/2021 - 1 | **5.1** |
| | | Influenza A/Cambodia/e826360/2020 - 1 | **1.7** |
| | | Influenza A/HongKong/45/2019 - 1 | **5.0** |
| | | Influenza B/Austria/359417/2021 - 1 | **3.5** |
| | | Influenza B/Washington/02/2019 - 1 | **2.0** |
| | | Influenza B/Phuket/3073/2013 - 1 | **5.8** |
| 5 | 700 | Influenza A/Michigan/45/2015 - 4 | **13.6** |
| | | Influenza A/Singapore/INFIMH-16-0019/2016 - 4 | **3.0** |
| | | Influenza B/Colorado/06/2017 - 4 | **8.6** |
| | | Influenza B/Phuket/3073/2013 - 4 | **13.6** |
| | | Influenza B/Phuket/3073/2013 – 3 | **4.3** |
| | | Influenza A/Wisconsin/67/2022 - 3 | **8.2** |
| | | Influenza A/Darwin/6/2021 - 2 | **4.2** |
| | | Influenza A/Darwin/6/2021 - 3 | **6.4** |
| | | Influenza B/Austria/359417/2021 - 2 | **4.8** |
| | | Influenza B/Austria/359417/2021 - 3 | **4.4** |
| 6 | 100 | Luciferase | **26.6** |
| | | Mini HA | **45.3** |
| | | Michi NA | **37.0** |
| | | Influenza A/Wisconsin/67/2022 -1 | **15.6** |
| | | Influenza A/Wisconsin/67/2022 -2 | **17.5** |
| | | Influenza A/Wisconsin/67/2022 -3 | **22.4** |
| | | Influenza A/Darwin/6/2021 -1 | **18.0** |
| | | Influenza A/Darwin/6/2021 -2 | **16.2** |
| | | Influenza A/Darwin/6/2021 -3 | **17.6** |
| | | Influenza B/Phuket/3073/2013 -1 | **19.1** |
| | | Influenza B/Phuket/3073/2013 -2 | **24.9** |
| | | Influenza B/Phuket/3073/2013 -3 | **13.7** |
| | | Influenza B/Austria/359417/2021 -1 | **14.8** |
| | | Influenza B/Austria/359417/2021 -2 | **16.0** |
| | | Influenza B/Austria/359417/2021 -3 | **13.2** |

*RNA constructs followed by a number indicate the different known codon optimization strategies applied for constructs.
