## Supplemental Table 2 for "Assay for Rapid Quantification of Capped and Tailed Intact mRNA"

**Supplementary Table 2. Accuracy and precision of two users on two different days in a 4-slide, 16-sample quantitative experiment**

| Expected % Capped | Precision (% RSD) | | |
|---|---|---|---|
| | User 1 | User 2 | Average |
| **15** | 2.5 | 3.7 | 3.1 |
| **20** | 7.2 | 1.1 | 4.1 |
| **25** | 7.3 | 1.0 | 4.2 |
| **30** | 4.8 | 2.7 | 3.7 |
| **35** | 3.0 | 1.7 | 2.4 |
| **40** | 1.7 | 1.3 | 1.5 |
| **45** | 2.4 | 1.3 | 1.9 |
| **50** | 3.6 | 3.0 | 3.3 |
| **55** | 4.6 | 2.5 | 3.6 |
| **60** | 4.3 | 3.0 | 3.7 |
| **65** | 3.4 | 2.9 | 3.2 |
| **70** | 3.4 | 2.5 | 2.9 |
| **75** | 4.3 | 3.2 | 3.7 |
| **80** | 3.7 | 3.2 | 3.4 |
| **85** | 4.7 | 2.8 | 3.7 |
| **90** | 3.6 | 2.5 | 3.1 |
| *Over all samples and operators (n=32)* | | | 3.2 ± 0.7 |

Supplier 2 mRNA tested by two operators (n=81 spots from n=4 replicates per sample).
