## Supplemental Table 3 for "Assay for Rapid Quantification of Capped and Tailed Intact mRNA"

**Supplementary Table 3. Expected % of capped and intact values for comparison between users and across two different days in a 2-slide, 2-sample quantitative experiment**

| | | **5’ Cap*Q* % Capped, Intact (LCL – UCL)*** | |
|---|---|---|---|
| | | **Day 1** | **Day 2** |
| Sample 1 (expected 51%) | User 1 | 48.8 (48.4 – 49.2) | 46.0 (45.6 – 46.4) |
| | User 2 | 51.9 (51.7 – 52.2) | 50.9 (50.5 – 51.2) |
| | User 3 | 57.2 (56.4 – 58.1) | 49.1 (48.4 – 49.8) |
| Sample 2 (expected 17%) | User 1 | 17.9 (17.7 – 18.2) | 15.5 (15.3 – 15.7) |
| | User 2 | 19.9 (19.7 – 20.0) | 16.5 (16.4 – 16.7) |
| | User 3 | 14.3 (14.1 – 14.6) | 14.9 (14.7 – 15.1) |

Supplier 2 mRNA tested by multiple operators over multiple days (n=81 spots from n=8 replicates per sample per experiment).

*The lower confidence limit (LCL) and upper confidence limit (UCL) represent the 95% confidence interval.
